# Functional and Clinical Implications of Extrachromosomal Circular DNA in the Human Germline

**DOI:** 10.1101/2024.06.02.597028

**Authors:** Melanie Evans, Shreya Rajachandran, Xin Zhang, Yanfeng Zhang, Karla Saner, Lin Xu, Kyle E. Orwig, Orhan Bukulmez, Haiqi Chen

## Abstract

Extrachromosomal circular DNA (eccDNA) originates from linear chromosomal DNA and can be found in various human cell types including the male germline. However, the functional effects and biogenesis mechanisms of the eccDNA in the human male germline are not well understood. Here, we developed a sequencing approach to extract eccDNA sequence information and the paired transcriptome information from the same cells. By applying this approach to human samples, we found evidence of transcriptional activities of germline eccDNAs. We also showed that patients with chronic diseases such as hypertension and diabetes had a significantly higher number of eccDNAs in the sperm than their healthy counterparts. This was, at least partly, due to an increased apoptosis signaling in the germline. Analysis of single cell RNA sequencing data of spermatogenic cells from diabetic patients *vs*. healthy individuals suggested that a dysregulation in the expression levels of multiple poly (ADP-ribose) polymerases may contribute to the increased amount of germline eccDNAs in diseased patients. In addition, we identified a potential horizontal transfer mechanism through which healthy sperm can take up eccDNAs from their surrounding microenvironment. Together, our results suggest that eccDNA may have functional effects on the germline, and it may serve as a non-invasive clinical biomarker for human health.

## Introduction

Prior work on cancer cells shows that deleted genomic DNA can self-circularize and exist as genetic fragments independent of the linear genome. These extrachromosomal DNAs - often referred to as ecDNAs - are of mega base pair long on average, derive from all parts of the genome, may contain genes that promote tumorigenesis, increase cellular heterogeneity to promote drug resistance, and can even re-integrate into the linear cancer genome to further disrupt genome integrity [1-6].

Interestingly, such circular DNAs were also found in normal somatic cells and germline cells [7-10] and were termed eccDNA (extrachromosomal circular DNA) to be distinct from the ecDNAs in cancer cells. Contrary to ecDNAs, however, the biological functions and biogenesis mechanisms of eccDNAs are less studied, especially in the context of the human germline.

In this study, we developed a method that enabled parallel sequencing of eccDNA and RNA in the same cells. Using this method, we discovered evidence of transcriptional activities of eccDNAs in the human male germ cells. We also showed that the number of sperm eccDNAs was significantly higher in patients with chronic diseases such as hypertension and diabetes than healthy men. This could be explained, at least partially, by an elevated level of apoptosis signaling in the germ cells. And the poly adenosine diphosphate ribose polymerases (PARPs) may be involved to mediate apoptosis-induced eccDNA generation. Finally, we discovered a potential new mechanism for healthy sperm to acquire exogenous eccDNAs.

## Results

### PEAR-seq detects eccDNA and RNA in the same cells

We developed <u>p</u>aired <u>e</u>ccDNA <u>a</u>nd <u>R</u>NA sequencing (PEAR-seq) by combining the current state-of-the-art eccDNA purification approach with an RNA-seq workflow (**Figure 1A**). To benchmark PEAR-seq, we used human sperm whose eccDNA content were previously characterized [8]. In PEAR-seq, total sperm DNA was separated from RNA through column-based purification. DNA was subjected to exonuclease digestion to enrich for circular DNA. A small subset of DNA was left undigested as a control. The DNA remaining after the digestion was fragmented and tagged with sequencing adaptors using Tn5 transposases followed by PCR amplification. This amplification approach overcomes the size bias of the rolling circle amplification (RCA)-based approach used in the previous circular DNA extraction protocols [7, 11] because smaller DNA circles are preferentially amplified than larger ones by RCA [10]. The paired RNA sample was subject to Illumina Total RNA Prep workflow to enable both coding and noncoding RNA quantification.

**Figure 1.**
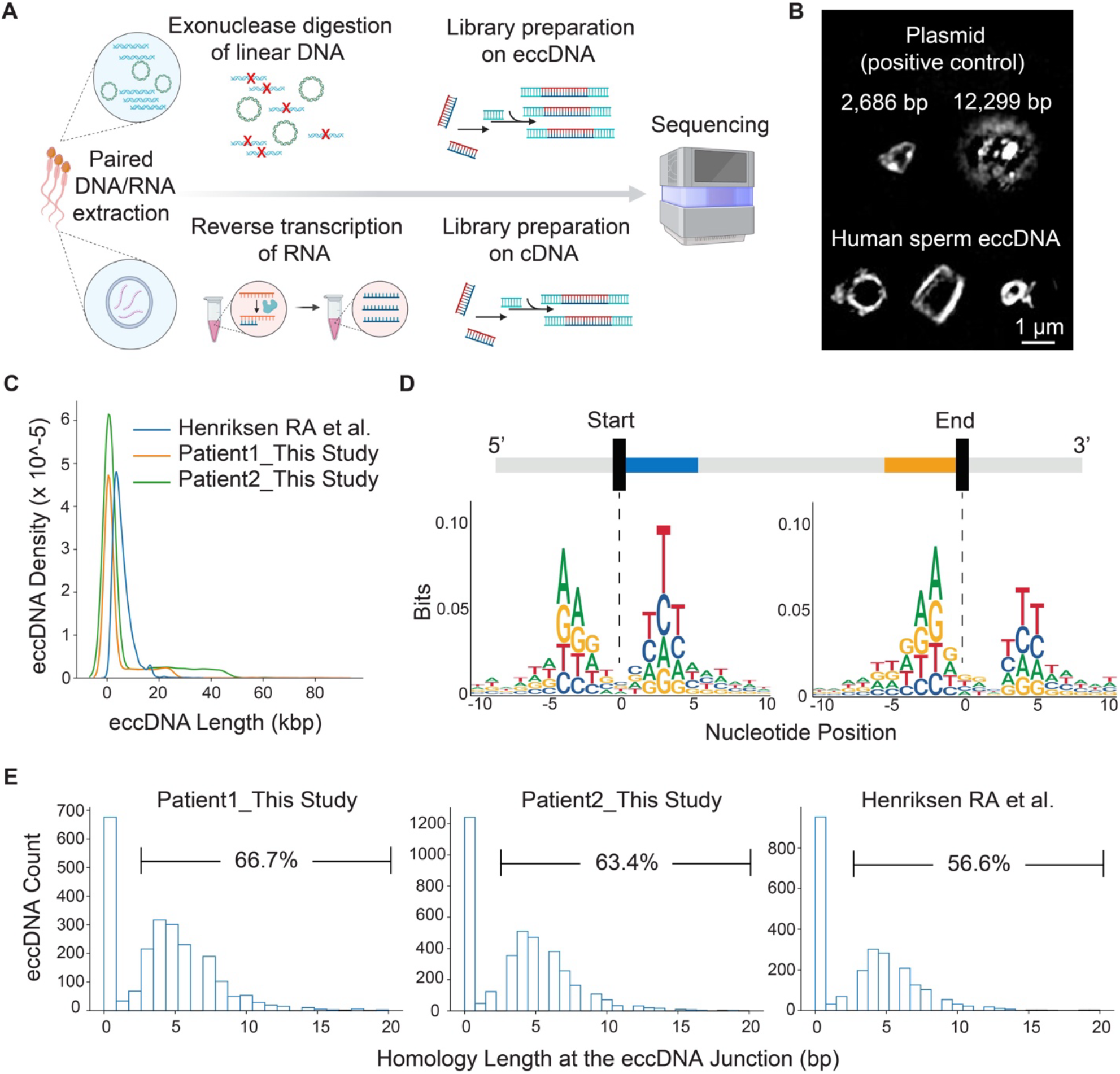
PEAR-seq identifies human sperm eccDNAs. (A) Schematic of the PEAR-seq workflow. Image was generated using BioRender. (B) SIM of plasmids and human sperm eccDNAs of varying sizes. (C) The size distribution of human sperm eccDNAs. (D) Trinucleotide motif sequences flanking the start and end positions of human sperm eccDNAs. (E) Distributions of the length of the homology sequences at the human sperm eccDNA junctions.

Sperm eccDNAs were extracted from two healthy patients using PEAR-seq. These eccDNAs were validated using the following orthogonal approaches. First, super resolution structured illumination microscopy (SIM) confirmed the circular structures of isolated sperm eccDNAs (**Figure 1B**); Second, by surveying the genome coordinates of the eccDNAs, we found that sperm eccDNAs derived from all parts of the human genome in a close to random manner (**Figure S1**) and varied in size from dozens of base pairs to hundreds of kilobase pairs (**Figure 1C**), consistent with the finding by Henriken *et al*. [8]; Third, by applying a permutation strategy, we showed that eccDNA-forming regions in the genome was depleted of long interspersed nuclear elements (LINEs) (enrichment score: 0.69; *p* value <10^-15) and was enriched, to a small extent, with short interspersed nuclear elements (SINEs) (enrichment score: 1.17; *p* value <10^-3). A similar observation was made in the eccDNAs isolated from mouse spermatogenetic cells [9]. Fourth, we identified enriched trinucleotide motif sequences flanking the start and end positions of sperm eccDNA molecules (**Figure 1D**). The same motifs were identified in eccDNAs isolated from the plasma of pregnant women [12] and human urine [13]. Finally, microhomology directed ligation has been indicated as one of the major mechanisms through which linear chromosomal DNA fragments are able to circularize to form eccDNAs [14]. Indeed, analysis of PEAR-seq-generated sperm eccDNA sequences and previously reported sperm eccDNA sequences showed that more than half of eccDNAs had ≥3bp homologous sequences flanking the start and end positions of eccDNA molecules (**Figure 1E**). Together, these data demonstrate that PEAR-seq can accurately identify eccDNAs in the human sperm.

### Human male germline eccDNAs show transcriptional activities

We next examined the DNA content carried by human sperm eccDNAs. On average, 19.1% of sperm eccDNAs contained at least one gene exon, and 4.92% of sperm eccDNAs carry at least one entire gene (**Figure S2**). Previous studies showed that ecDNAs in cancer cells can transcribe the oncogenes they carry [1, 3]. Thus, we set out to leverage PEAR-seq to test if male germline eccDNA can drive the transcription of its DNA content. In a bulk population, gene transcripts coming from eccDNAs are mostly indistinguishable from the products of the same gene on the linear genome. To circumvent this challenge, we devised two approaches to look for hybrid gene products that would be unique to eccDNAs if transcription did occur on eccDNAs.

In the first approach, we sifted through the coordinates of human sperm eccDNAs identified by PEAR-seq and looked for eccDNA loci in the genome whose start and end points, once connected via circularization, would generate a hybrid gene that is made of exons from two different genes. We then performed PCR using the paired RNA sample as a template to enrich for potential transcripts from the hybrid gene. The existence of the hybrid gene transcript was further confirmed by next generation sequencing. **Figure 2** shows two examples of the eccDNA transcription events observed from a patient sperm sample using this approach. In the first example, we found that an eccDNA generated by the circularization of a 24 kb region of chromosome 19 resulted in a hybrid gene that was made of the first exon of *ZNF439* and the last 3 exons of *ZNF440* (**Figure 2A**). High throughput sequencing of the PCR product generated using the paired RNA as the template confirmed the existence of the chimeric transcript spanning the corresponding eccDNA junction (**Figure 2B**). In the second example, circularization of another 22 kb region on chromosome 19 lead to an eccDNA which contained a hybrid gene consisted of the first 7 exons of *ZNF615* and the last exon of *ZNF 614* (**Figure 2C**). The existence of this hybrid transcript was also confirmed by high throughput sequencing of the PCR product (**Figure 2D**).

**Figure 2.**
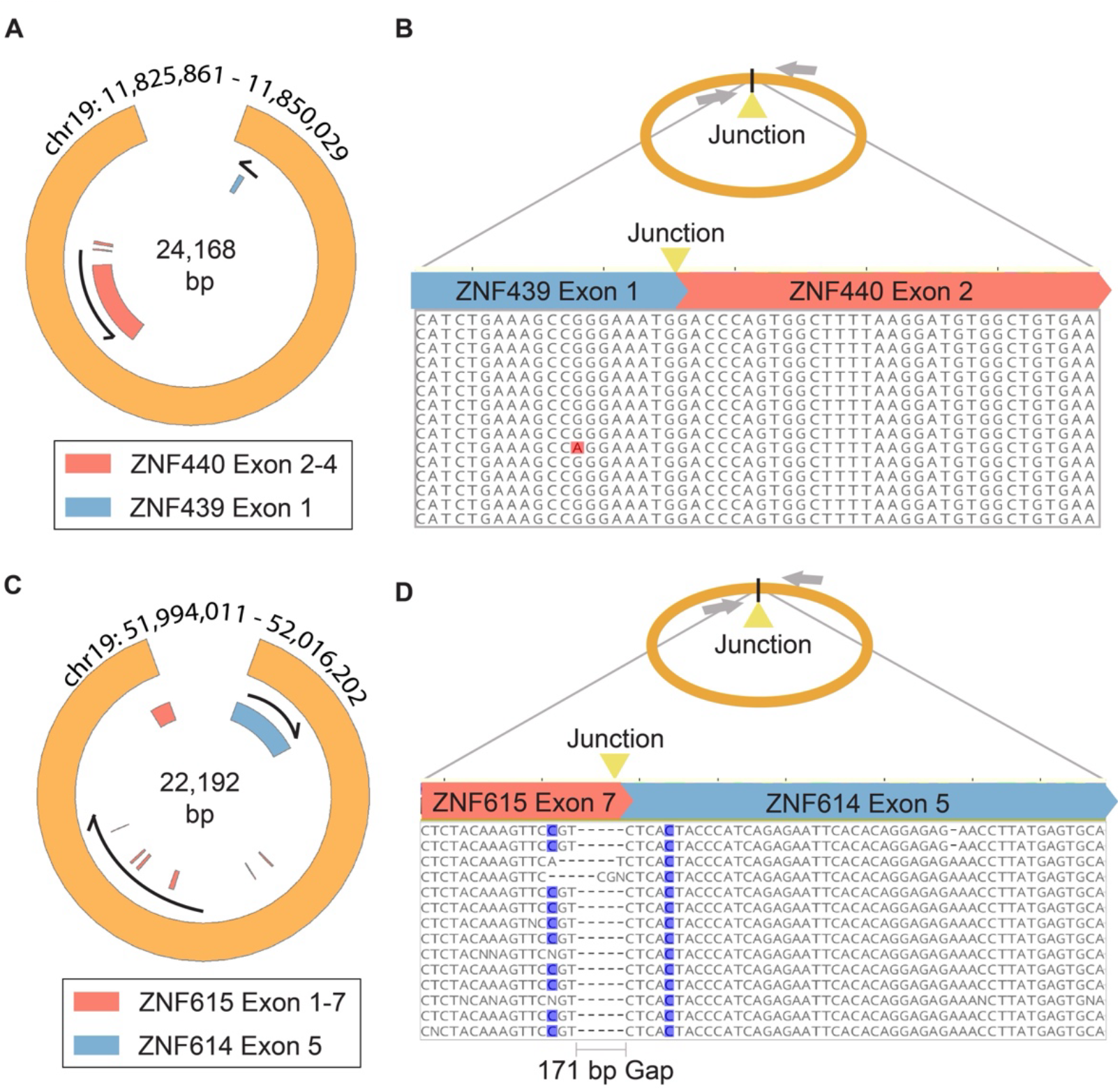
Examples of germline eccDNA transcription events. (A) Schematic of the eccDNA containing the first exon of *ZNF439* and the last 3 exons of *ZNF440*. (B) Representative sequencing reads from the paired RNA sample generated using PEAR-seq aligned to the junction of the eccDNA. Red block indicates sequencing error in the reads. (C) Schematic of the eccDNA containing the first 7 exons of *ZNF615* and the last exon of *ZNF 614*. (D) Representative sequencing reads from the paired RNA sample generated using PEAR-seq aligned to the junction of the eccDNA. In this case, the reads contain a 171 gap when compared to the reference eccDNA sequence, likely due to splicing of the hybrid mRNA. Blue blocks indicate SNPs in the patient genome.

In the second approach, we sequenced the paired eccDNA-RNA libraries generated from sperm samples from two healthy individuals using PEAR-seq. We systematically characterized split-transcript reads from the RNA libraries, defined as reads in which their left part aligned to the end region and the right part to the start region of an eccDNA (Methods). Using this approach, we identified 29 transcription events in one sample, and 114 events in the other sample (**Table S1**).

Together, our data suggested that at least a subset of male germline eccDNAs can be transcribed. Since transcriptional activities are largely silenced in sperm, the eccDNA-generated transcripts we have detected in mature sperm were likely produced during earlier stages of spermatogenesis.

### The amount of sperm eccDNA is associated with individual health status

We next examined the clinical relevance of germline eccDNA by comparing the amount of sperm eccDNAs from healthy individuals with those from patients diagnosed with hypertension or diabetes. To avoid age being a confounding factor, only samples from individuals between age 30 and 55 were used. We found that the number of sperm eccDNAs was significantly elevated in patients with hypertension and patients with diabetes when compared with healthy men (**Figure 3A**).

**Figure 3.**
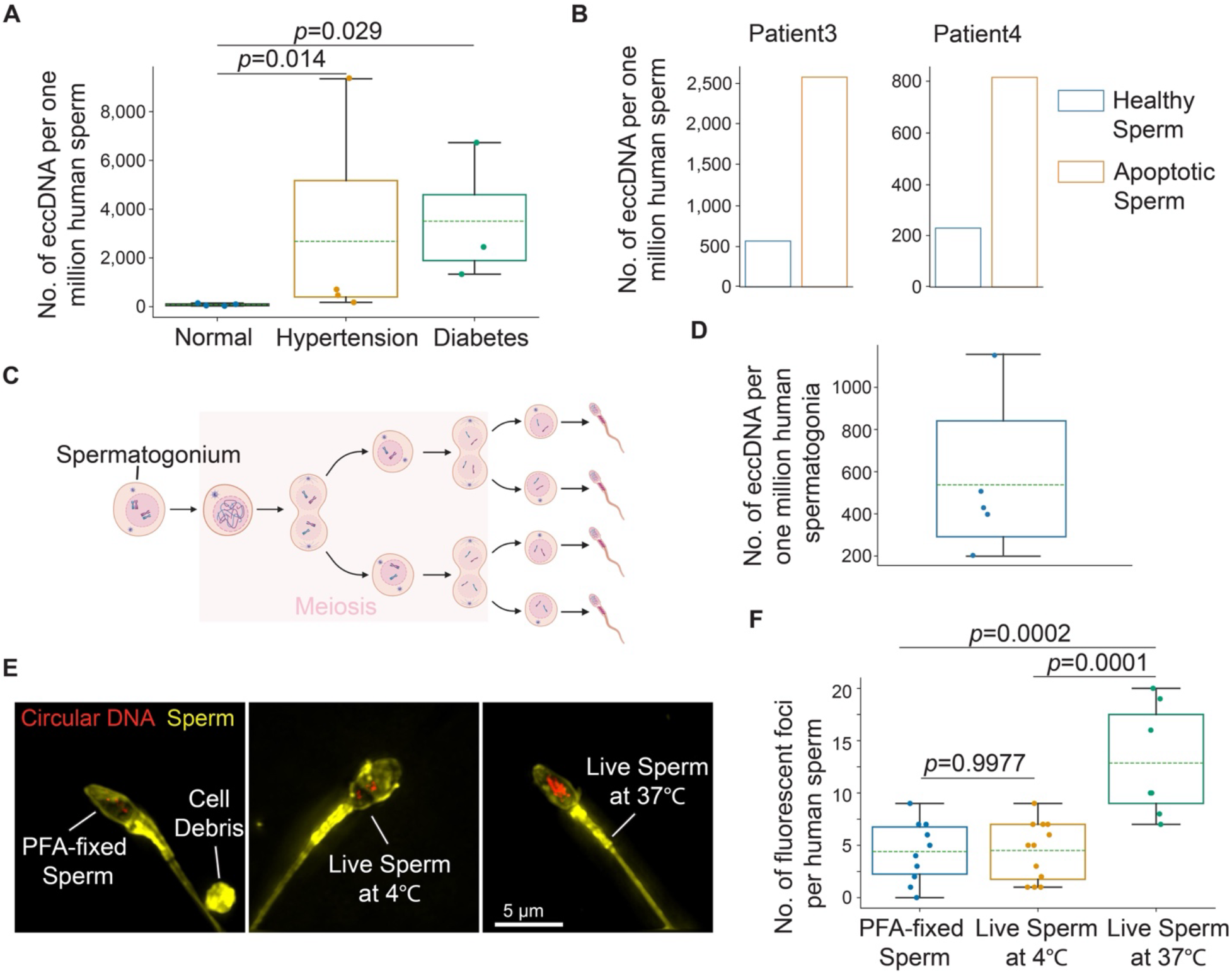
Mechanisms of eccDNA biogenesis in the human male germline. (A) The number of sperm eccDNAs in patients with different health status. *p* values were calculated using the Wilcoxon test. (B) The number of sperm eccDNAs in apoptotic sperm *vs*. healthy sperm from the same semen sample. (C) Schematic of human spermatogenesis. Image was generated using BioRender. (D) The number of sperm eccDNAs in human spermatogonia. (E) SIM images of human sperm (yellow) containing circular DNA (red) under each condition. (F) Quantification of the number of circular DNA loci in the human sperm under each condition. *p* values were calculated using one-way ANOVA with post-hoc Tukey HSD test.

### Apoptotic signaling promotes male germline eccDNA formation

We then asked why diseased patients had more sperm eccDNAs than healthy individuals. Previous work showed that there was a significant increase in sperm DNA damage in hypertensive men [15]; and that the apoptosis signaling was significantly increased in sperm from males with diabetes [16]. Of interest, recent studies suggested that cellular apoptosis can lead to eccDNA generation [17]. Thus, we hypothesized that apoptosis in the germ cells was one of the mechanisms underlying the disease-induced increase in the amount of sperm eccDNAs.

To test this hypothesis, we first analyzed single cell RNA sequencing (scRNA-seq) data of testis samples from normal and diabetic patients [18, 19]. GO term analysis showed that the apoptotic signaling pathways was indeed one of the significantly enriched signaling pathways in spermatogenic cells of diabetic patients (adjusted *p* value <0.05) (**Figure S3A**). To examine if apoptosis induced eccDNA generation in the male germline, we isolated apoptotic sperm and healthy sperm from the same semen sample using flow cytometry (Methods) and quantified their respective eccDNA content. Consistent across two healthy individuals, we found that apoptotic sperm contained approximately 4-fold more eccDNAs than healthy live sperm (**Figure 3B**). Together, out data suggested that patients with chronic diseases such as diabetes had elevated apoptotic signaling in the germline, leading to an increase in germ cell DNA damage and the generation of more eccDNAs than healthy individuals.

To further identify the key players involved in the disease-induced germline eccDNA formation, we surveyed the differentially expressed genes (DEGs) (adjusted *p* value < 0.05) in the spermatogenetic cells from diabetic patients *vs*. healthy individuals using the scRNA-seq data. Intriguingly, the expression of several poly (ADP-ribose) polymerases (PARPs) was significantly altered in the spermatogenic cells of diabetic patients (**Figure S3B**). PARPs were reported to show DNA repair activities when DNA strand breaks occur in the male germ cells due to oxidative stress, chromatin remodeling, or cell death [20]. Thus, the disruption in the expression of PARPs in the germline of diabetic patients may contribute to the biogenesis of eccDNAs.

### Horizontal transfer as a potential mechanism of eccDNA acquirement in the human male germ cells

When we compared the eccDNA quantity of healthy sperm with that of the apoptotic sperm from the same semen sample, we found that healthy sperm contained eccDNA, albeit at a much lesser extent than the apoptotic sperm (**Figure 3B**). This suggested that male germ cells can acquire eccDNAs through other mechanisms besides the intrinsic apoptotic signaling pathway. Indeed, meiotic recombination has been suggested to contribute to eccDNA biogenesis in healthy germ cells [8]. To test if meiosis was the main stage during which healthy germ cells acquired eccDNAs, we set out to quantify the eccDNA content in pre-meiotic spermatogonia (**Figure 3C**). Of interest, we found that healthy human spermatogonia contained eccDNAs at a level comparable to that of the healthy human sperm (**Figure 3D**). This indicated that meiotic recombination might not be the only mechanism by which eccDNA was generated in healthy germ cells. We thus set out to examine other possibilities of eccDNA acquirement by healthy germ cells.

Apoptosis can often result in the release of the DNA content out of the apoptotic cell [21]. Because we have shown that eccDNAs can be generated in apoptotic sperm, we asked if eccDNAs released from apoptotic sperm can be taken up by healthy sperm. To this end, we co-incubated fluorescently labelled circular DNA with PFA-fixed human sperm (i.e., dead sperm) at 37 °C, live human sperm at 4 °C, and live human sperm at 37 °C, respectively (Methods). By imaging these samples using SIM, we showed that live sperm at 37 °C acquired significantly more circular DNA than PFA-fixed sperm and live sperm at 4 °C (**Figure 3E&3F**). This observation was further validated by extracting eccDNAs from these sperm samples using PEAR-seq and quantifying the acquired circular DNA using qPCR (**Figure S4**). Together, these results showed that human sperm can actively take up cell-free eccDNAs in their surrounding microenvironment likely through an energy-dependent endocytosis mechanism. This observation supported the possibility of the horizontal transfer of eccDNAs from apoptotic sperm to healthy sperm.

## Discussion

The extent to which eccDNAs occur in the male human germline has been vigorously characterized [8]. In this study, we provided new insights into the functions and biogenesis of germline eccDNAs.

Our first observation was that male germline eccDNAs can be transcribed. From two independent patient samples, we found a total of 143 transcripts across eccDNA junctions (**Table S1**), suggesting that at least a fraction of eccDNAs resides inside the nucleus. The number of identified eccDNA transcripts is likely an underestimation of the true number. This is because some eccDNA-derived transcripts may not cover the eccDNA junctions, which would not be detected using our current approaches. Indeed, we found that an average of 4.92% of sperm eccDNAs carry at least one entire gene (**Figure S2**). These gene products, if transcribed from eccDNAs, would be indistinguishable from those of the same genes on the linear genome. Since we detected eccDNA-derived transcripts in mature human sperm and transcriptional activities are largely shut down during spermiogenesis, these eccDNA-derived transcripts were likely to be generated before spermiogenesis and were stored in sperm without being degraded. In cancer cells, ecDNAs with copies of proto-oncogenes, such as *cMYC* and *EGFR*, were shown to promote tumorigenesis via elevating the protein level of these genes [2, 3]. Thus, it would be interesting to examine if germline eccDNA-derived transcripts can be translated and if the resulted proteins exert roles in sperm functions and/or embryo development after fertilization.

Our second observation was that the amount of sperm eccDNAs was inversely correlated with a male’s health status as patients with hypertension or diabetes had more sperm eccDNAs than healthy individuals. This observation will allow us to explore the potential applications of sperm eccDNAs as noninvasive biomarkers for health monitoring. To this end, a larger sample size with samples from patients with diverse genetic backgrounds and ages will be needed.

We also investigated the mechanism(s) by which chronic diseases promoted the generation of sperm eccDNAs. First, we demonstrated that microhomologous sequences (≥3bp) were present at boundaries of more than half of all the sperm eccDNAs (**Figure 1E**), suggesting that the microhomology-mediated end joining was one of the major mechanisms for circularizing DNA fragments to form germline eccDNAs. Next, we showed that sperm undergoing apoptosis contained much more eccDNAs than healthy sperm from the same semen sample (**Figure 3B**). This suggested that human germline eccDNAs might be the products of apoptosis. Finally, by analyzing scRNA-seq data of testicular cells from diabetic men *vs*. health men, we showed that the apoptotic signaling pathway was perturbed in the diabetic germ cells and that PARPs might play a key role in mediating apoptosis-induced germline eccDNA formation (**Figure S3**).

The exact mechanism(s) through which healthy human sperm acquire eccDNAs remain to be determined. Our work indicated that meiotic recombination might not be the only source of eccDNAs in healthy germ cells since healthy spermatogonia which have not entered meiosis also contained eccDNAs (**Figure 3D**). This raised an interesting question regarding the biogenesis of eccDNAs in healthy germ cells as germline genome integrity is closely maintained in a healthy germline, making genome deletions much less unlikely compared with apoptotic germ cells. A previous study showed that when co-incubating live mouse sperm with plasmids, mouse sperm were able to actively take up these plasmids [22]. We made a similar observation that live human sperm could acquire artificial circular DNA when incubated together at 37 °C. This was likely mediated through an energy-dependent endocytosis process rather than passive diffusion because both dead human sperm and live sperm incubated at 4 °C contained significantly less circular DNA. This led to a possibility that healthy sperm could acquire eccDNAs that were released into the microenvironment from apoptotic cells, resulting in the gain of additional genetic materials by the healthy sperm. Thus, it would be interesting to examine if the human seminal fluid contains eccDNA. In addition, it remains to be determined if eccDNAs carried by healthy sperm can be transmitted to the next generation.

### Methods Human Samples

Adult human sperm samples were obtained from the fertility clinic at UT Southwestern Medical Center. At the fertility clinic, couples who underwent in vitro fertilization (IVF) were required to leave semen samples cryopreserved at the clinic in case they were unable to give a fresh specimen on the day of IVF. This has resulted in a large collection of banked backup semen samples from patients with various health status at the clinic. According to standard clinical regulations, these backup semen samples should be discarded after the successful completion of the IVF cycles following patient consent. An IRB-approved protocol allowed us to gain access to these unused backup semen samples for basic research purposes. Information about the deidentified sperm samples used in this study was listed in **Table S2**.

Adult human testicular samples were from 3 healthy men (donor #1: 18-28 years old, black; donor #2: 40-50 years old, white; donor #3: 41 years old, race unknown); Samples were obtained through the University of Pittsburgh Health Sciences Tissue Bank and Center for Organ Recovery (CORID No. 686). All samples were de-identified.

### PEAR-seq

Cryopreserved human semen samples were thawed, and sperm were washed with PBS before subject to the Qiagen Allprep DNA/RNA Mini Prep kit (Cat. #: 80204) following the manufacturer’s instructions to collect both the DNA and RNA contents from the same patient sample. Plasmid Safe ATP-dependent DNase kit (BioSearch Technologies; Cat. #: IDE3110K) was used to digest chromosomal linear DNA. For 2 μg of total sperm DNA, 30 units (3 μL) of exonuclease, 4 μL of ATP (25 mM), and 10 μL of 10x reaction buffer were added to reach a reaction volume of 100 μL. The digestion was performed at 37°C for at least 3 days. Each day an additional 4 μL of ATP (25 mM), 0.6 μL of 10x reaction buffer, and 3 μL of exonuclease were added into the reaction mix to continue the enzymatic digestion reaction. The exonuclease treated sample was used to confirm elimination of chromosomal linear DNA by quantitative polymerase chain reaction (qPCR), using chromosomal markers such as *ACT1* (5’-TCCGTCTGGATTGGTGGTTCTA-3’ and 5’-TGGACCACTTTCGTCGTATTC-3’) and *COX5B* (5’-GGGCACCATTTTCCTTGATCAT-3’ and 5’-AGTCGCCTGCTCTTCATCAG-3’). Each 20 μL of qPCR reaction contains 2 μL of exonuclease-treated sample, primers at a final concentration of 150 nM, 0.2 μL of 100x SYBR Green dye (ThermoFisher Scientific; Cat. #: S7563), and 10 μL of Q5 Hot Start High-Fidelity 2X Master Mix (NEB; Cat. #: S7563). 1 ng of undigested total sperm DNA from the same sample was used as control. The following qPCR reaction condition was used: 30 sec at 98°C, followed by 35 cycles of 10 sec at 98°C, 15 sec at 65°C, and 30 sec at 72°C. If qPCR result indicated the existence of remaining undigested linear DNA, an additional day of exonuclease digestion was performed. If the result confirmed the elimination of linear DNA, the exonuclease in the digestion solution was heat inactivated by incubating the solution at 70 °C for 30 minutes.

The digested DNA was purified using a NucleoSpin Gel and PCR Clean-Up Kit (Takara; Cat. #: 740609), and then quantified using a Qubit dsDNA Quantification Assay Kit (ThermoFisher Scientific; Cat. #: Q32851). A Nextera XT Library Prep Kit (Illumina; Cat. #: FC-131-1024) was used to generate a sequencing library following the manufacturer’s instructions except that DNA at 0.4 ng/μL instead of 0.2 ng/μL was used as input. For the paired RNA sample, an Illumina TruSeq Stranded Total RNA Library Prep Gold Kit (Cat. #: 20020598) was used to generate an rRNA-depleted total RNA sequencing library (including both the coding and non-coding RNAs). The concentration and size distribution of the sequencing libraries were quantified using the Agilent D1000 High Sensitivity Screentape Kit (Cat. #: 5067-5584) kit on an Agilent TapeStation 2200 system. Libraries were sequenced on an Illumina NextSeq 2000 system at the UT Southwestern Medical Center McDermott Center Next-generation Sequencing Core and an Illumina NovaSeq X Plus system at Novogene Corporation with a sequencing run cycle of 150 bp/8 bp/8 bp/150 bp (read 1/index 1/index 2/read 2).

### Identification of eccDNAs with Circle-Map

The adaptor sequences of the paired end reads were trimmed using BBDuk under BBMap v38.46. The trimmed reads were aligned to the hg38 version of the human genome using BWA MEM v.0.7.5. All downstream BAM/SAM file processing analyses were performed using Samtools v1.6. Circle-Map v.1.1.4 was used to detect eccDNAs by first extracting all soft-clipped, hard-clipped, and discordant reads to a new BAM file using the ‘ReadExtractor’ command and then executing the ‘realignment’ module while keeping all parameters at their default settings. Mitochondrial DNA was removed from downstream analysis.

### eccDNA Imaging

eccDNAs in 10 mM Tris buffer (pH 8) were mixed with YOYO-1 Iodide (ThermoFisher Scientific; Cat. #: Y3601) at a 1:10 dye-to-base pair ratio. The sample was placed in a 35 mm glass-bottom (MatTek Corporation; Cat. #: P35G1.514C), mounted with prolong diamond antifade mountant (ThermoFisher Scientific; Cat. #: P36961), and sealed with a cover glass. Images were obtained on a DeltaVision OMX SR Super-resolution Microscope using an Olympus 60x 1.42 NA Oil Immersion PSF objective. Plasmids of known sizes were used as positive controls.

### Identification of eccDNA Junctional Motifs

To explore the motif patterns flanking the sperm eccDNA junctions, we scanned the base compositions from 10 bp upstream to 10 bp downstream of the start and end positions for each eccDNA coordinate. Homer v4.9 was used to calculate the nucleotide frequency within the 21 bp windows. ggseqlogo v0.2 was used to plot the frequency. A random set of coordinates were generated to match the length and chromosome distribution of the eccDNA coordinates. The same analysis was performed on this random set.

### Detection of eccDNA Transcription Events

Segemehl v0.3.4 was used in split-read mode, without realignment, to detect split-transcript reads from the total RNA-seq data generated using PEAR-seq. Split-transcript reads were defined as reads in which the 5’ part of the reads aligned to the 3’-end of a genome locus and the 3’ part of reads to the 5’-end of the locus. Split-transcript read intervals that entirely overlapped with the detected eccDNA coordinates were collected.

### Flow Sorting of Human Sperm

Semen samples from healthy males were thawed. Depending on the sperm count, cells were split equally into several microcentrifuge tubes with each tube containing approximately 500,000 sperm. Cells were centrifuged at 500 g at 4°C for 5 min. Supernatant was discarded, and each pellet was resuspended in 500 uL of 1X Annexin V Binding Buffer (BD Biosciences; Cat. #: BDB556454). The cells were then recombined and counted with a phase contrast hemocytometer (Hausser Scientific; 3200).

To prepare cells for flow cytometry, the following antibody and dye were used. Annexin V-PE antibody (BD Biosciences; Cat. #: BDB556422) was diluted in 1X Annexin V Binding Buffer at a 1:20 dilution. The reagents from the live/dead fixable violet dead cell stain kit (ThermoFisher Scientific; Cat. #: L34964) was diluted in 1X Annexin V Binding Buffer at a 1:1000 dilution. Three aliquots with 1 million cells each were prepared as control groups: 1) a negative control group in which cells were not incubated with any antibody or dye; 2) a live/dead stain only group in which cells were incubated with the diluted live/dead violet stain dye at the room temperature for 20 min; and 3) a Annexin V-PE only group in which cells were incubated with the diluted Annexin V-PE antibody at the room temperature for 20 min. The treatment group contained the rest of the sperm incubated with both the diluted live/dead violet stain dye and diluted Annexin V-PE antibody at the room temperature for 20 min. All groups were protected from light during the incubation.

After incubation, cells were centrifuged at 300 g at 4°C for 5 minutes and resuspended in 1X Annexin V Binding Buffer to a final concentration of 2 million cells/mL. Flow cytometry was performed on a FACS LSRFortessa cell sorter (BD Biosciences). The three control groups described above were used for calibration and gating. In the treatment group, sperm positive for Annexin V-PE were sorted into the apoptotic group and cells negative for both live/dead violet stain and Annexin V-PE were sorted into the healthy group.

### Isolation of Human Spermatogonia

Cryopreserved human testicular tissues from three healthy adult donors were thawed quickly in a 37°C water bath and washed in Hank’s Balanced Salt Solution (HBSS) (ThermoFisher Scientific; Cat. #: 24020141). Tissues were then digested sequentially with 2 mg/mL Collagenase IV (Worthington Biochemical; Cat. #: LS004188), 0.25% Trypsin-EDTA (ThermoFisher Scientific; Cat. #: 25200114), and 3.5 mg/mL DNase I (Worthington Biochemical; Cat. #: LS002147) in HBSS at 37°C. The cell suspension obtained was filtered through a 40 µm cell strainer (Falcon; Cat. #: 352340), and dead cells were removed through magnetic activated cell sorting (MACS) using the dead cell removal kit (Milteny Biotec; Cat. #: 130-090-101). MACS-sorted live cells were stained with PE rat anti-human CD49f/ITGA6 (10 µL/10^6^ cells; BD Biosciences; Cat. #: 555736) in 1X DPBS supplemented with 1% FBS (ThermoFisher Scientific; Cat. #: 10082-147), then incubated in 1X DPBS containing anti-PE MicroBeads (2 µL/10^6^ cells; Milteny Biotec; Cat. #: 130-048-801). The ITGA6-positive cells were collected as spermatogonia.

### Analysis of Human Testis scRNA-seq Data

The scRNA-seq data for human normal testis was obtained from Gene Expression Omnibus (GEO) through accession number GSE106487. The testis scRNA-seq data from diabetic patients was obtained from Song K. *et al*. [19]. DEGs between normal and diabetic patients in each spermatogenic cell type was calculated using the Seurat v4 FindMarker function. The Wilcoxon test was used to calculate *p* values. Genes with an average log-transformed difference greater than 0.5 and an adjusted *p* value less than 0.05 were defined as DEGs. GO term enrichment analysis was performed using clusterProfiler v4.11.1.

### Circular DNA Acquirement Assay

YoYo-1 Iodide was incubated with 2.5 ug of circular DNA at a 1:12.5 dye to base pair ratio in 1X PBS at the room temperature for 30 min. To remove excess dye, DNA was run through a QIAprep Spin Miniprep Kit Column (Qiagen; Cat. #: 27106) and eluted in 1X PBS.

Semen samples from healthy males were thawed and then divided equally into three groups: 1) a 37 °C group, 2) a 4°C group, and 3) a PFA-treated group. Each group contained approximately 500,000 sperm. Cells were spun down at 500 g at 4°C for 5 min and the supernatant was discarded. For the 37 °C and 4 °C groups, the pellets were resuspended in sperm washing medium (Irvine Scientific; Cat. #: 9983). For the PFA-treated group, cells were resuspended in 4% PFA (ThermoFisher Scientific; Cat. #: J19943.K2). All three groups were incubated at the room temperature for 10 min, spun down, and resuspended in 500 uL of sperm washing medium. Equivalent volumes of YoYo-1-stained circular DNA (approximately 500 ng of DNA per group) were then added to each sample group. Cells in the 37 °C group and the PFA-treated group were placed in a cell culture incubator with 5% CO_2_ at 37 °C for 1 hr whereas cells in the 4 °C group were incubated at 4 °C for 1 hr. After incubation, sperm were spun down and washed twice with 0.1% dextran sulfate (Milipore Sigma; Cat. #: S4030) in the sperm washing medium to eliminate residual DNA on the surface of sperm. Samples were then resuspended in 500 uL of sperm washing medium.

To prepare samples for imaging, 50 uL of sperm from each group were incubated with CellMask deep red plasma membrane stain (ThermoFisher Scientific; Cat. #: C10046) at a 1:1000 dilution in a cell culture incubator with 5% CO_2_ at 37 °C for 10 min. Samples were then washed with 1XPBS and fixed in 1% PFA for 5 min at the room temperature. Samples were then spun down and resuspended in 50 uL of 1X PBS. 8 uL of samples were added onto a 35 mm glass bottom dish. Samples were mounted with prolong diamond antifade mountant and sealed with a cover glass. Imaging was done with done on the DeltaVision OMX SR Super-resolution Microscope using an Olympus 60x 1.42 NA Oil Immersion PSF objective.

## Data Availability

The raw sequencing data supporting the findings of this study are available in the NCBI BioProject database with BioProject ID PRJNA1117395.

## Code Availability

Custom code is available at https://github.com/HaiqiChenLab/human_sperm_eccDNA.

## Acknowledgments

We thank Sihan Wu for helpful discussions and Dorothy Mundy for providing assistant with SIM.

H.C. acknowledges support from the Cecil H. and Ida Green Center for Reproductive Biology Sciences Endowment.

## Author Contributions

H.C. conceived and supervised the project. M.E., S.R., and X.Z performed experiments. H.C., M.E., S.R., X.Z., and Y.Z. performed data analysis. K.E.O. provided human testicular samples.

K.S. and O.B. provided human semen samples. K.S., L.X., O.B. and K.E.O provided consultations. H.C., M.E., and S.R. wrote the manuscript with input from all authors.

**Figure S1.**
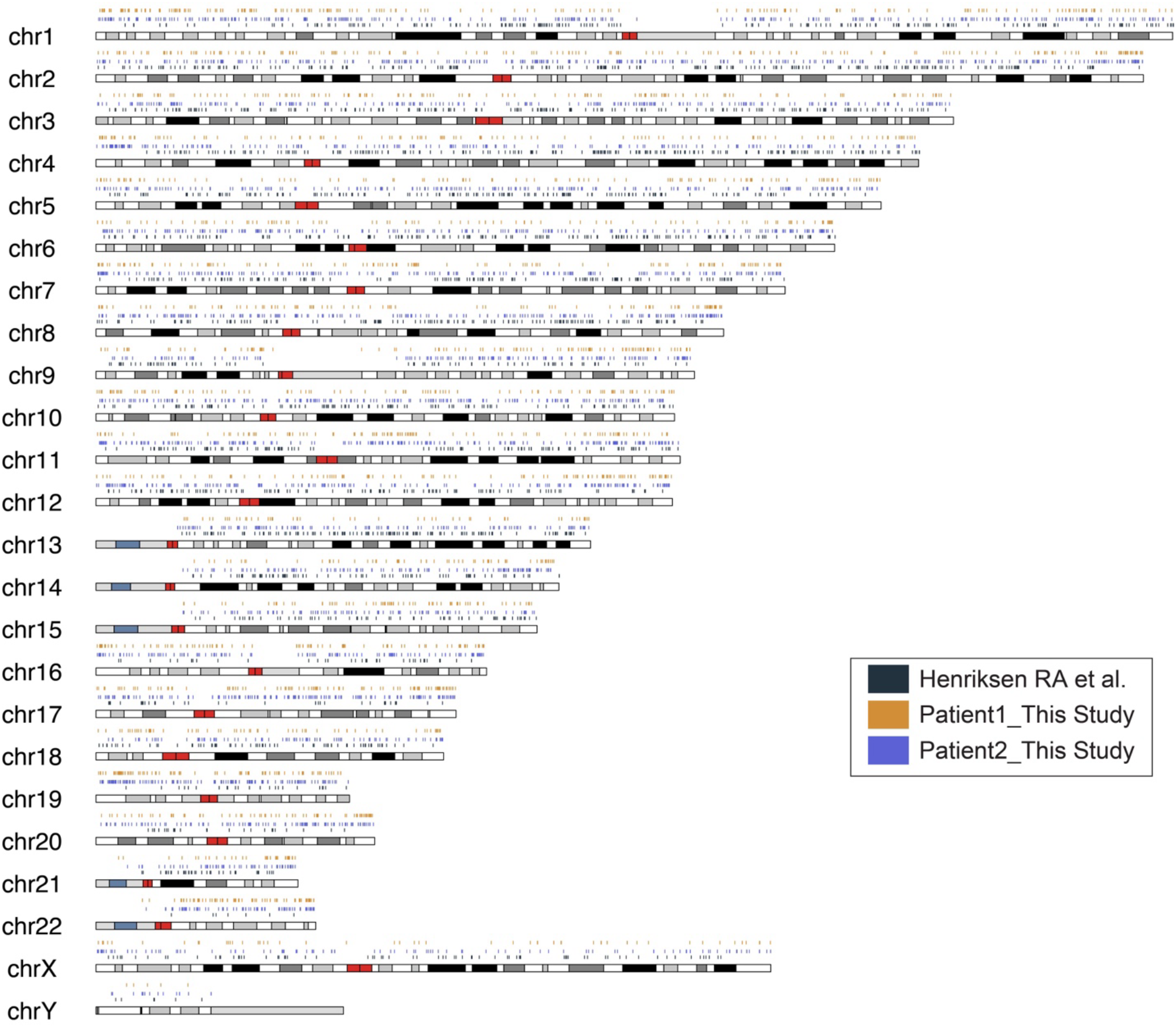
Human sperm eccDNAs derive from every chromosome. Origins of eccDNAs in the human genome in three independent samples. Colored lines above the chromosomes represent eccDNAs.

**Figure S2.**
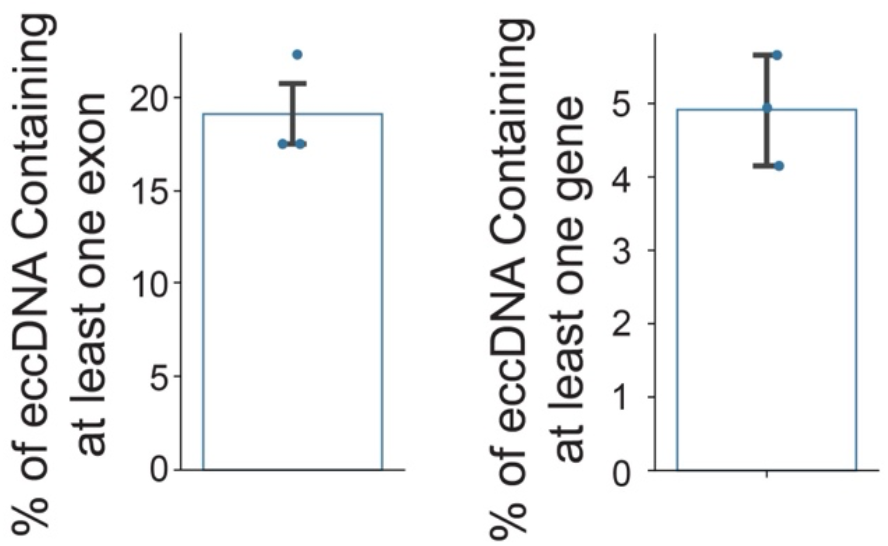
Human sperm eccDNAs contain exons and whole genes. Bar plots showing the percentage of human sperm eccDNAs containing at least one gene exon and at least one entire gene.

**Figure S3.**
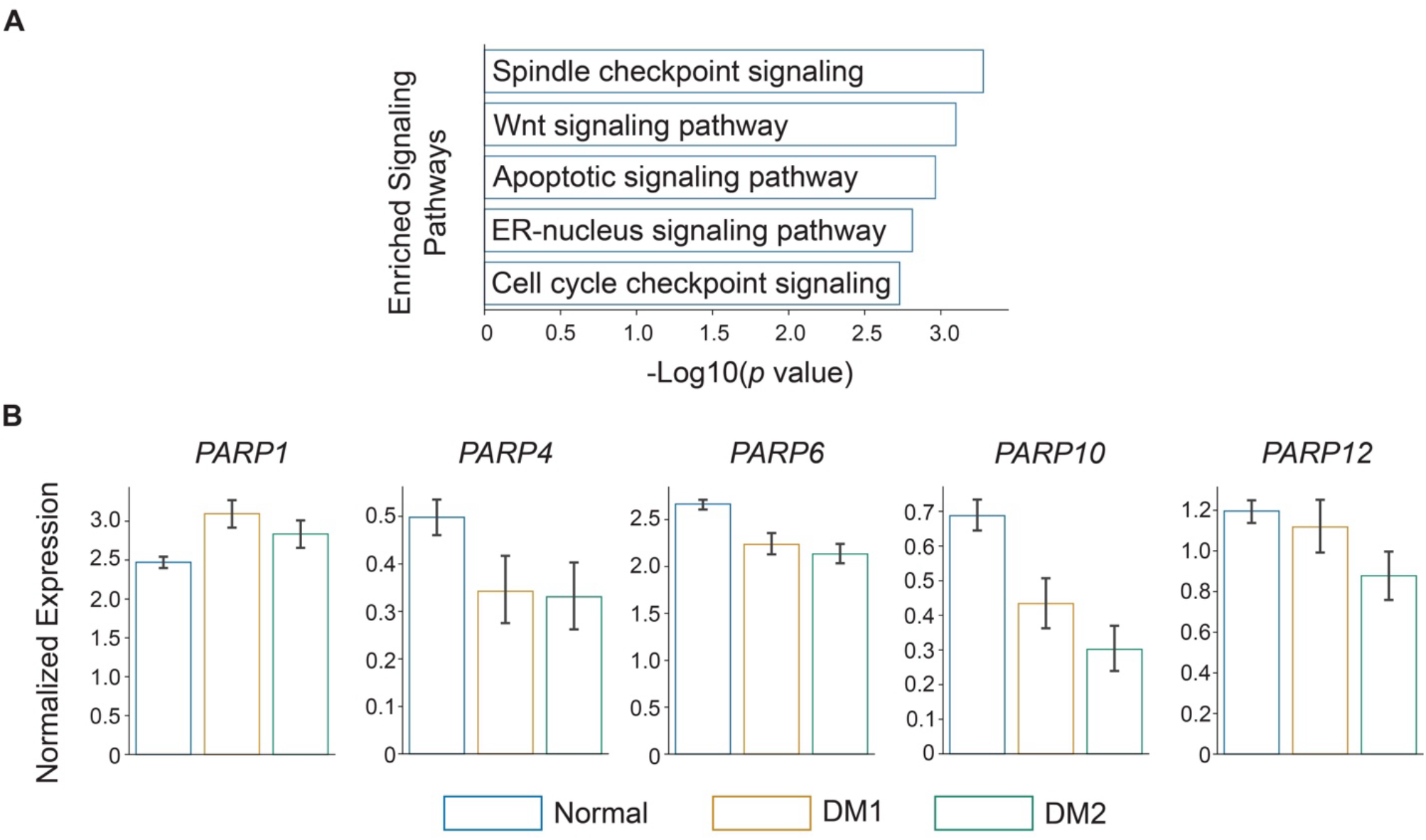
The apoptotic signaling was significantly altered in the germline of diabetic males. (A) Representative enriched GO terms in the DEGs of spermatogenic cells from diabetic *vs*. normal patients. *p* values were calculated using a hypergeometric test. (B) The expression level of several *PARP* genes in spermatogenic cells from normal and diabetic (DM1 and DM2) patients.

**Figure S4.**
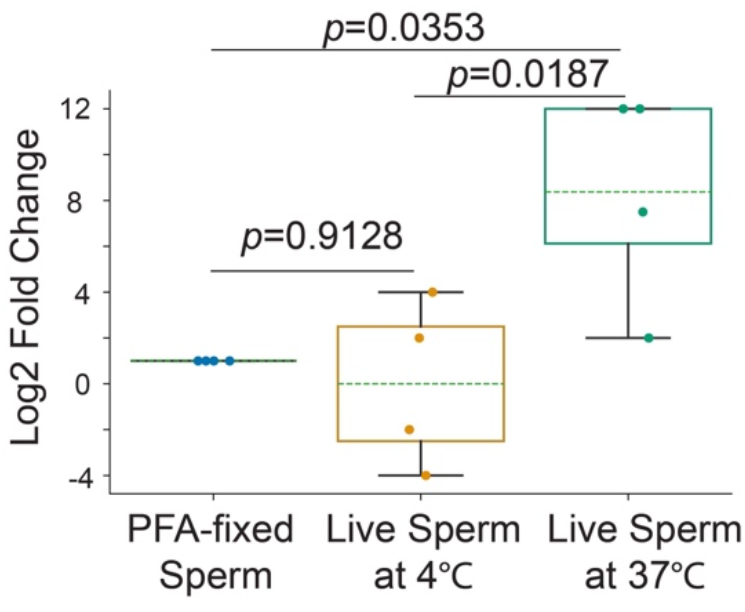
Live sperm at 37 °C acquired significantly more circular DNA from the surrounding environment than sperm from other groups. qPCR quantification of the amount of circular DNA acquired by the human sperm under each condition. *P* values were calculated using one-way ANOVA with post-hoc Tukey HSD test.

## Notes

### Competing Interest Statement

The authors have declared no competing interest.

